# Multiple origins of endogenous virophage and polinton-like virus in the halophilic protist *Halocafeteria seosinensis*

**DOI:** 10.64898/2026.07.28.741278

**Authors:** Ronie Haro, L. Gallot-Lavallée, Cedric Blais, John M. Archibald

## Abstract

Virophages and Polinton-like Viruses (PLVs) are related viral elements classified within the supergroup *Polisuviricotina*. While virophages typically depend on large dsDNA viruses (*Nucleocytoviricota*) for replication in eukaryotic hosts, the replication strategies of most PLVs are unknown. While both virophages and PLVs can exist as stand-alone entities or be integrated into host genomes, their co-occurrence within a single eukaryotic genome is rare. We investigated the chromosome-scale nuclear genome assembly of the halophilic protist *Halocafeteria seosinensis* and discovered 41 endogenous PLVs and 36 virophage sequences, most of which are full-length elements; together they comprise 6.2% of the genome. These viral elements belong to six PLV and seven virophage subtypes. Notably, we found genes shared between *H. seosinensis* PLVs and virophages, suggesting active genetic exchange between them. We also observed supraparasitism, with MULE DNA transposons and LINE retrotransposons frequently embedded within the viral genomes. Our study reveals dynamic interactions between viral elements and host mobile DNA in a halophilic protist, expanding our understanding of viral diversity in extreme environments.

**SIGNIFICANCE:** Virophages and polinton-like viruses (PLVs) are enigmatic viral elements found in many aquatic habitats. Our understanding of their diversity stems mainly from analysis of environmental sequence data, but they have also been found integrated into the genomes of cultured protists. The evolutionary impacts of such integrations are poorly understood. In this study, we reveal the co-occurrence of multiple PLV and virophage subtypes that together constitute a substantial fraction of the genome of the halophilic protist *Halocafeteria seosinensis*. These distantly related viral lineages coexist within the same genome and share a common gene pool, forming chimeric arrangements with host transposons and with one another. The *H. seosinensis* genome thus serves as a dynamic arena for viral gene exchange and genome remodelling, reshaping host genome architecture and potentially conferring immunity to giant viruses.

## INTRODUCTION

Eukaryotic nuclear genomes are often rich in transposable elements (TEs), including endogenous viral elements (EVEs), which can impact genome architecture and evolution (1, 2). Among EVEs, polintons (or Mavericks) are a prominent class of large, self-replicating eukaryotic DNA mobile elements initially discovered in animal genomes and originally classified as transposons (3). Polintons are ∼20 kilobase pairs (Kbp) in length and bound by Terminal Inverted Repeats (TIRs). A hallmark of polintons is the presence of genes for a protein-primed type B DNA Polymerase (pPolB) and a retroviral-like Integrase (RVE) (from which the name is derived), as well as DNA-packaging ATPase (ATPase) and maturation protease (PRO) genes. Although not initially described, most polintons also encode Major Capsid Protein (MCP) and minor Capsid Protein (mCP) homologs (4) typical of double-stranded DNA (dsDNA) viruses, suggesting that they can form virions (as yet not observed) in addition to having a transposon-like lifestyle.

Virophages are small (20-30 Kbp) dsDNA viruses (5–7) whose capsid, ATPase, and PRO proteins are homologous to those of polintons and other dsDNA viruses. A subset of virophages also possesses a Polinton-like pPolB and RVE, which speaks to their evolutionary relatedness to polintons (8, 9). Virophages were discovered serendipitously (9–12) when they were isolated alongside large dsDNA viruses of the phylum *Nucleocytovirocota* (NCLDVs), which they require for their replication within a eukaryotic host. Some virophages confer anti-viral protection to the hosts (9, 10, 13, 14) and thus may significantly impact the population dynamics of eukaryotes and the DNA viruses that infect them (15–17). While virophages were initially isolated as virions, endogenous virophage genomes were later identified in several eukaryotic genomes (18–20), suggesting that at least some of them have an integrated stage. Notably, the heterotrophic protist *Cafeteria burkhardae*, which belongs to the protist phylum Bigyra, harbours a variety of endogenous virophages (21) whose replication is activated by the presence of a replicating dsDNA virus called CroV (14).

Polinton-like Viruses (PLVs) are mobile elements recently discovered in metagenomic data from aquatic environments (22, 23). Their MCP, mCP, and ATPase genes are distantly related to those of polintons and virophages. Only a small fraction of PLVs also encode a PRO (24). The replication and integration apparatus of PLVs is heterogeneous (22, 24–27) and can involve genes homologous to those of polintons (pPolB and RVE) (22, 24, 26) or virophages (22, 25, 27). Some PLVs lack an obvious mCP gene (28) and others have no detectable integration gene (25). PLVs are comprised of several clades with highly divergent capsid proteins (24). PLV genomes can exist as stand-alone molecules (22), as nuclear genome integrants (20, 25), or associated with *Nucleocytoviricota*-type particles (29). Recently, the PLV Gezel-14T was shown to produce virions when co-infecting the brown alga *Phaeocystis globosa* alongside *P. globosa* virus 14T (PgV-14T) (30), suggesting that some PLVs rely on giant viruses for their replication, similar to virophages. In the case of the PLV of the green alga *Tetraselmis striata*, the *T. striata* virus (TsV-N1) (31) only requires the host to release virions and is therefore considered an ‘autonomous’ virus. The replication strategies of most PLVs remain unclear.

Based on genome size and overlapping gene content, polintons, PLVs and virophages were recently proposed to constitute separate classes within the viral supergroup *Polisuviricotina* (32). While a few studies have shed light on their replication strategies (31, 33), the range of ‘lifestyles’ exhibited by the members of this supergroup is unclear. Moreover, a growing body of literature has documented a shared gene pool among polintons, PLVs, and virophages. The size of the pool and the tempo and mode(s) of genetic exchange among the three *Polisuviricotina* classes remain to be determined.

Although there are exceptions (18, 19, 21, 25, 34), most PLV and virophage sequences detected in eukaryotic genome assemblies reside on short contigs (see (20) and references therein), which makes it difficult to determine their genomic contexts and coding capacities. We and others (20) detected sequences related to virophages and PLVs in a short-read genome assembly of the halophilic protist *Halocafeteria seosinensis* (35), a relative of the virophage-containing *C. burkhardae* (21). Using Oxford Nanopore long-read DNA sequencing, we recently produced a chromosome-scale assembly for *H. seosinensis* (36). Here, we show that endogenous PLVs and virophages are a prominent feature of the *H. seosinensis* genome and have exchanged genes with one another over recent evolutionary timescales.

## RESULTS

### The *H. seosinensis* genome is enriched in virophages and PLVs

We screened our chromosome-level assembly of *H. seosinensis* (36) for endogenous virophages and PLVs using structural genes from virophages (MCP, mCP, ATPase, PRO) and PLVs (MCP, mCP, ATPase) as queries (HMM and BLASTp searches). Candidate regions were further examined for terminal inverted repeats (TIRs) and GC-content shifts and were classified as virophages or PLVs based on MCP sequence similarity networks (SSN) built from published virophage and PLV sequences. We identified 41 complete PLVs and 36 complete virophages across 43 of the 60 pseudochromosomes, together accounting for 6.2% of the nuclear genome (Table S1). These elements were frequently situated close to telomeric repeats, and complete or partial viral genomes were identified at 13 pseudochromosome ends (Fig. 1A, Fig. S7a). In some cases, multiple virophages were arranged in tandem, and viral regions were frequently associated with other mobile elements (e.g., LINE-1 retrotransposons, Helitrons and MULEs; see below). Notably, with one exception (Fig. S4J), only virophage genomes were detected in subtelomeric regions.

**Fig. 1.**
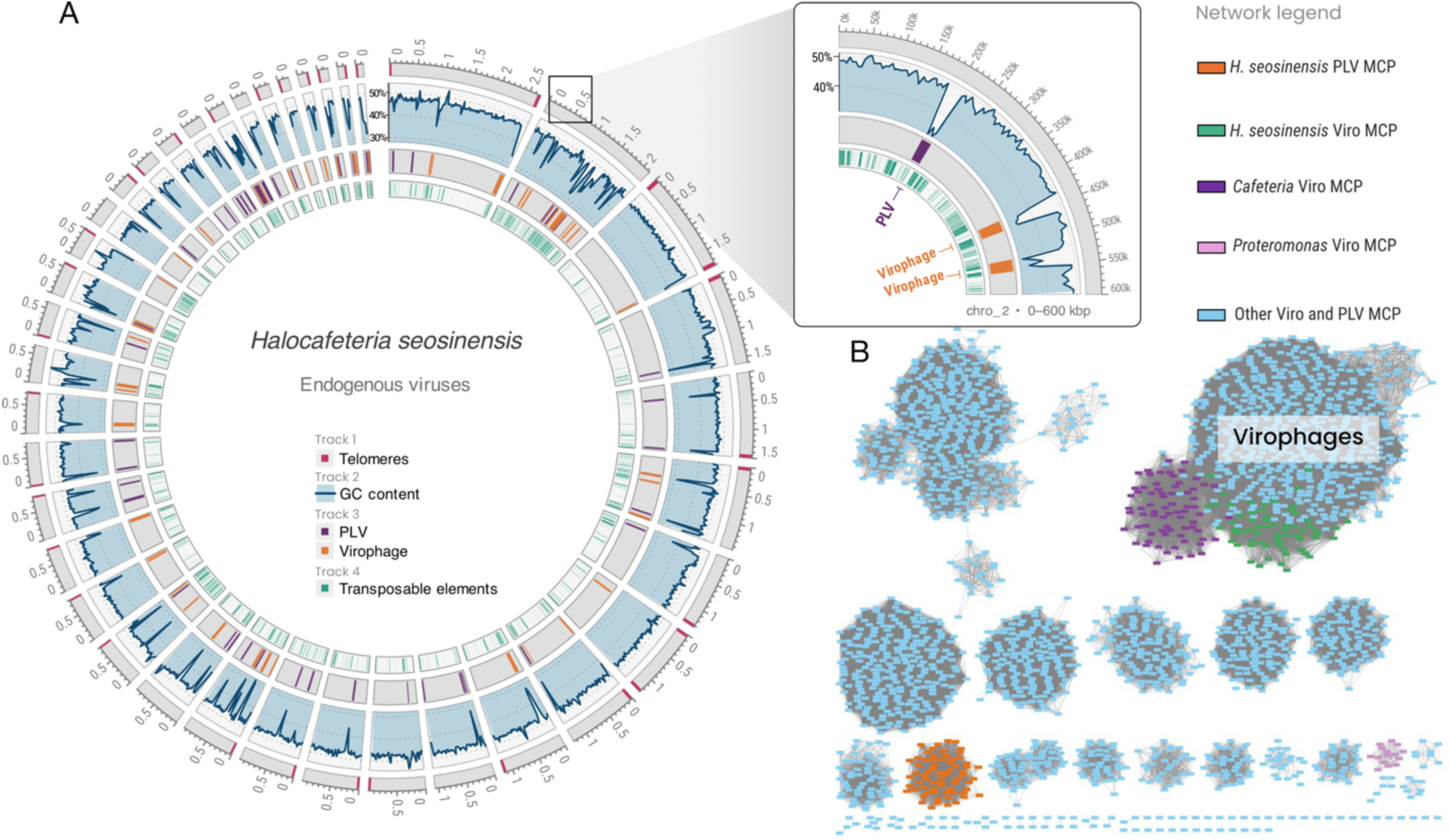
Diversity of endogenous viruses in the 39 Mbp nuclear genome of *Halocafeteria seosinensis*. **A.** Genomic assembly of *H. seosinensis* with annotated virophages and PLVs. The inset box shows a 600 Kbp genomic region of chromosome 2 highlighting specific insertions and their distinct GC contents. **B**. Major Capsid Protein (MCP) sequence similarity network of virophages and PLVs. Virophage MCPs form a distinct large cluster grouping *H. seosinensis* (green), *Cafeteria burkhardae* (purple) and other virophages obtained from metagenomic studies (light blue). PLV MCPs are distributed across multiple clusters; *H. seosinensis* PLVs form a unique stand-alone cluster (orange). MCPs associated with the stramenopile *Proteromonas* are in purple.

Complete virophages are slightly larger on average than PLVs (24,556 bp versus 23,580 bp) (Table 1, Table S1), and their TIRs are longer (808 bp versus 485 bp). Both classes have GC contents (35% for virophages and 34% for PLVs) lower than the host genome in which they reside (47%), making many of them easy to recognize (Fig. 1A). Most complete elements contain the expected core genes, including structural, replication, and integration genes, and are flanked by TIRs. Some PLVs (n=5) and virophages (n=12) contain a full complement of core genes but lack obvious TIRs, suggesting either sequence degeneration or alternative boundary structures. We also detected fragmented virophage (n=18) and PLV (n=12) sequences lacking most core genes (Table 1, Table S1); they likely represent decaying remnants or elements that depend on proteins encoded by complete copies. Some still retain TIRs and target side duplications (TSDs) (Table 1, Table S1), which could indicate transposition/integration after loss of structural genes. Those with replication and integration genes might have adopted a transposon-like lifestyle.

**Table 1.** Summary of *Halocafeteria seosinensis* endogenous virophages and polinton-like viruses (PLVs)

|  | Complete<br>elements | Average<br>size (bp) | Average<br>TIR size<br>(bp) | Average<br>TIR %<br>identity | TSDs<br>detected | Partial<br>elements<br>(most core<br>genes<br>present) | Elements<br>with<br>few/non-<br>core<br>genes<br>(with TIRs) |
| --- | --- | --- | --- | --- | --- | --- | --- |
| PLVs | 41 | 23,580 | 485 | 98.3 | 31 | 5 | 8 (6) |
| Virophages | 36 | 24,556 | 808 | 98.3 | 26 | 12 | 18 (7) |

### *H. seosinensis* PLVs and virophages belong to distinct subclades

Endogenous virophages and PLVs in *H. seosinensis* form host-specific groups that are not closely related to elements from other Bigyra, including *Cafeteria burkhardae* (Bicosoecida), which harbors only endogenous virophages (21) (Fig. 1B). This co-occurrence of PLVs and virophages has been observed in a few other unicellular eukaryotes (20) including the chlorarachniophyte alga *Bigelowiella natans* (18), but in *H. seosinensis* the PLV-like MCPs form a unique cluster distinct from known PLVs, including those in *Proteromonas lacertae* (Fig. 1B). This contrasts with earlier surveys in which *H. seosinensis* PLVs appeared to cluster with other Bigyra (20); this likely reflects the use of different similarity thresholds for detecting endogenous viruses across unicellular eukaryotic genomes. In our study, *H. seosinensis* virophage MCPs group together as a cohesive unit (Fig. 1B) and form a well-supported monophyletic clade (Fig. S1A) that is not specifically related to *C. burkhardae* virophages — despite the close phylogenetic relationship between their hosts (36, 37) — or any other virophages. A similar pattern is seen in mCP phylogenies (Fig. S1B).

Structural-gene phylogenies reveal substantial intragenomic diversity within these host-specific endogenous viruses. Based on MCP trees (Fig. 2A,B), we defined six PLV subtypes and seven virophage subtypes, and these subtype groupings are consistently recovered in trees for mCP (Fig. S1B–D) and ATPase (Fig. S2A, B). PRO-based phylogenies could be inferred only for virophages, because *H. seosinensis* PLVs lack PRO (Fig. 2C, Fig. S2C). The one notable exception is virophage subtype 7, which clusters along with subtype 5 virophages according to ATPase and PRO phylogenies (Fig. S2A, C). This pattern could reflect either phylogenetic artifacts or genuine exchange of replication-related genes between these subtypes.

**Fig. 2.**
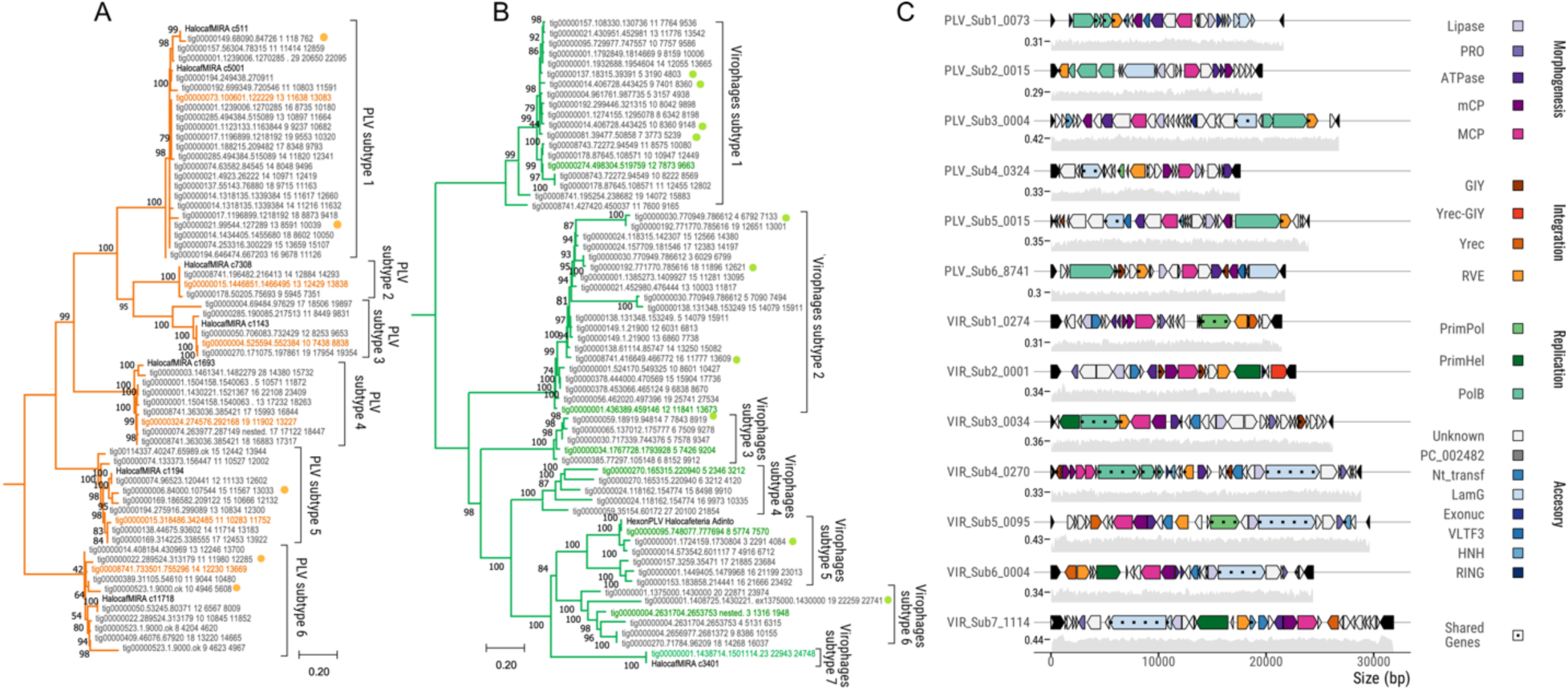
*H. seosinensis* PLVs and virophages belong to multiple subtypes. **A**. Maximum likelihood phylogenetic tree of PLV MCPs showing the subtree formed by *H. seosinensis* sequences. The tree was inferred using the LG+F+R7 model and 1000 ultrafast bootstrap replicates; it is based on 562 sequences aligned with MAFFT-LINSI, retaining 381 sites with <30% gaps (full tree reconstruction not shown). Orange labels indicate MCPs of PLVs included in panel C, while light orange dots denote MCPs contained in partial genomes. Bold labels represent MCPs reported in a previous study (20). **B**. Maximum likelihood phylogenetic tree of virophage MCPs. The alignment included 629 sequences and 510 sites (<30% gaps). The tree was constructed under the LG+F+R10 model with 1000 ultrafast bootstrap replicates. The subtree containing all *H. seosinensis* virophage MCPs is shown here (a broader phylogenetic reconstruction is shown in Fig. S1A). Green labels indicate MCPs of virophages represented in panel C; green dots indicate homologs from partial genomes. Bold labels denote MCPs characterized in a previous study (20, 22, 23). The scale bars show the inferred number of amino acid substitutions per site. **C**. Genome map of PLV and virophage subtype representatives from *H. seosinensis*. Colour-coded genes represent conserved functions across viral elements. Genes are grouped into hypothetical functional modules: morphogenesis, integration, replication, and accessory. Dotted patterns indicate shared genes. Gray graphs below each element illustrate fluctuations in GC content (500 bp sliding window). Terminal inverted repeats (TIR) are displayed as black arrows. Viral genome representations were generated using gggenomes (https://github.com/thackl/gggenomes).

Compositional and synteny analyses support a history of repeated acquisition and diversification. Most PLVs and virophages in *H. seosinensis* have GC contents lower than their host pseudochromosomes (Fig. 1A, Fig. 2C), but PLV subtype 3 (42%) virophage subtypes 5 and 7 (44%) have GC averages closer to the host (48%) (Table S3); this is attributed to amelioration following an ancient integration event. Within these subtypes, some elements retain lower GC, suggesting more recent acquisitions. Subtype-specific genome alignments and collinearity analyses (e.g., subtype 1; Fig. S8) suggest that certain viral genomes underwent relatively recent whole-genome duplications, whereas other members of the same subtype show extensive rearrangements (Fig. S8). Taken together with distinct protein-cluster profiles for each subtype, these patterns suggest that *H. seosinensis* has captured a diverse set of PLVs and virophages through multiple independent events, followed by subtype-specific expansion and remodelling within the host genome.

### Overlapping gene repertoires and mobile replication modules

The gene-sharing network from Vcontact2 (Fig. 3A) shows that endogenous virophages in *H. seosinensis* are strongly connected to other virophages (1,568 edges shared with 247 other virophages), reflecting the high conservation of their structural genes (37). In contrast, PLVs have fewer links to other PLVs (only 20 edges shared with 2 other PLVs), consistent with their greater sequence diversity (20, 24). Unexpectedly, 124 shared edges exist between the virophages (n=27) and PLVs (n=28) (Fig. 3A, Table S4), indicating extensive sharing of protein clusters beyond their structural modules (Fig. 1B, Fig. S1 and S2). Some clusters, such as RVE integrases (PC_000012), protein-primed DNA polymerases (pPolB) (PC_000001, PC_000010, PC_000053), RING finger proteins (PC_000452), primase–polymerases (PC_002058), and a peptide release factor (PC_002482), occur in most members of several PLV and virophage subtypes (Table S4). By contrast, HNH endonucleases (PC_000097), GIY nucleases (PC_000087), and LamG-domain (PC_001637) are present in only subsets of elements across multiple subtypes (Table S4). Phylogenetic analyses of these proteins (Fig. 3B and C, Fig. S3) support two main origin scenarios. In the first scenario, the trees suggest that the gene exchanges happened before or shortly after the initial spread of a virus subtype within the *H. seosinensis* genome, either pre-or post-integration. In the second, exchange appears to have occurred after multiple viruses of the same subtype were established. Genes of the first type often include core replication and integration functions, while those in the second involve accessory or mobility-related roles.

**Fig. 3.**
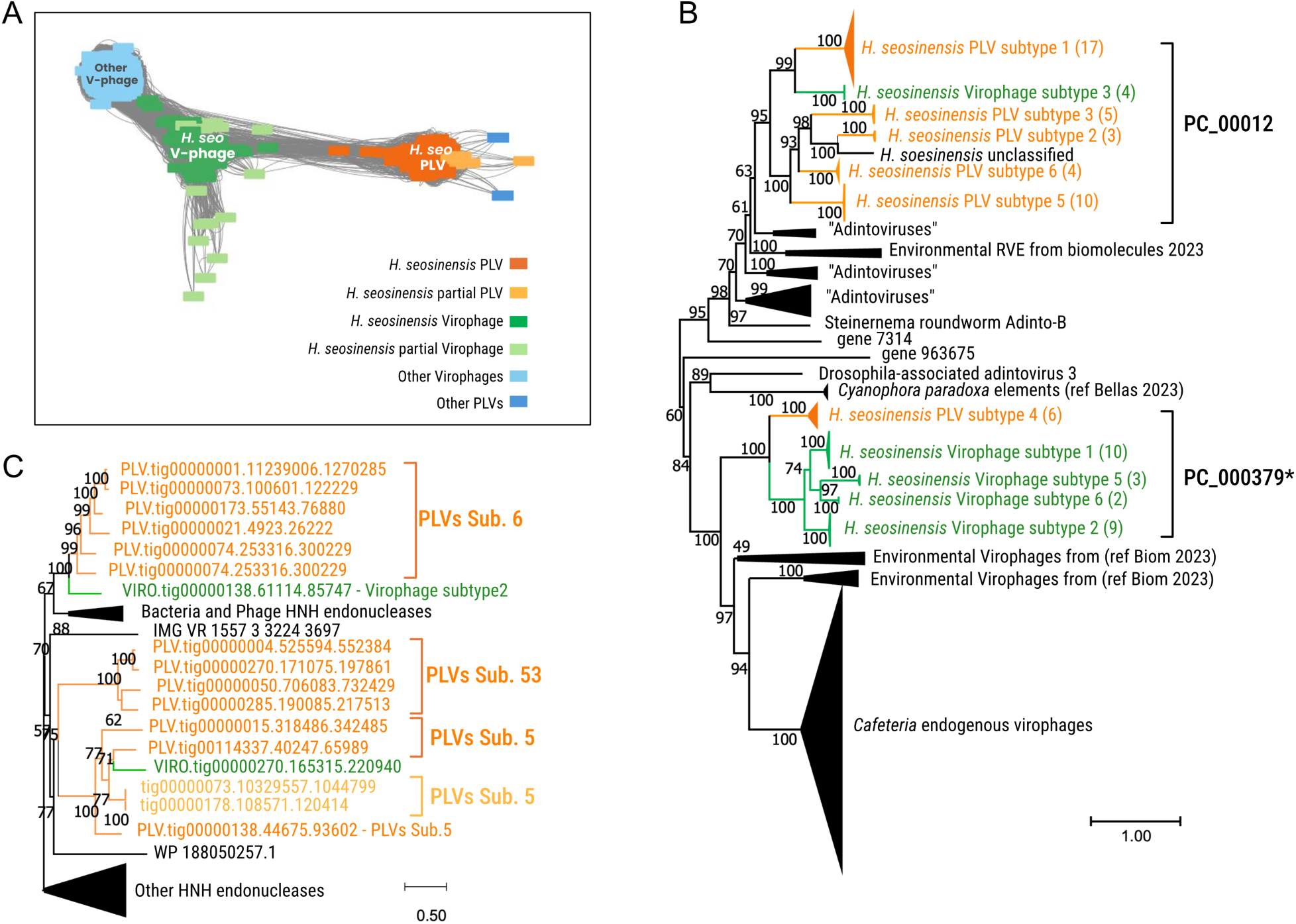
Gene exchange between *H. seosinensis* endogenous virophages and PLVs. **A**. Vcontact2 network of *H. seosinensis* endogenous virophages and PLVs proteomes, including other *Polusviricotina*. *H. seosinensis* endogenous virophages and PLVs are tightly clustered, suggesting shared gene content. The network was visualized with Cytoscape using an edge-weighted spring-embedded layout. **B**. Maximum likelihood phylogenetic tree of RVE integrases inferred under the LG+F+R6 model and 1000 ultrafast bootstrap replicates. A subset of sequences from PC_000012, PC_000379 and PC_000056 was selected with UCLUST (3). The final alignment of 229 sequences contains 346 sites with <40% gaps. Sequences from virophage subtypes 4 and 7 were excluded due to insufficient length. The single PLV subtype 4 sequence from PC_000056 clusters within PC_000379 (highlighted by a *). **C**. Phylogenetic tree of HNH endonucleases (PC_000097) inferred under the VT+F+I+G4 model and 1000 ultrafast bootstrap. 38 sequences were aligned with MAFFT, and 157 sites with <30% gaps were retained. No orthologs were found in *H. seosinensis* outside of the endogenous viruses. Scalebar shows inferred amino acid substitutions per site.

In stark contrast to the monophyly of the structural proteins (i.e., MCP, mCP, ATPase, PRO), replication and integration genes in *H. seosinensis* endogenous elements are highly diverse and appear to move between PLVs and virophages. Across subtypes, these elements encode multiple distinct replication proteins (n=5) and integration proteins (n=3) (Fig. 4A), often located near one end of the viral genome (Fig. 4C), as expected under non-homologous gene replacement (26). Some virophages even carry two different replication genes (e.g., pPolB together with a D5 primase–helicase, or D5 primase–helicase together with PolA-S3H), suggesting transient states in ongoing replacement. A subset of virophages, but not PLVs, also encode both RVE integrase and a Tyrosine recombinase integrase (Yrec) (Fig. 2C and 4A), a combination that has been linked to *C. burkardae* integrated virophages (21) and giant virus genomes (22, 38). This may also be true for *H. seosinensis* virophages, in which one integrase mediates integration into the eukaryotic host genome and the other into a giant virus. Together, these features point to a highly mobile replication/integration toolkit that is shared and reshuffled between PLVs and virophages within the same host.

**Fig. 4.**
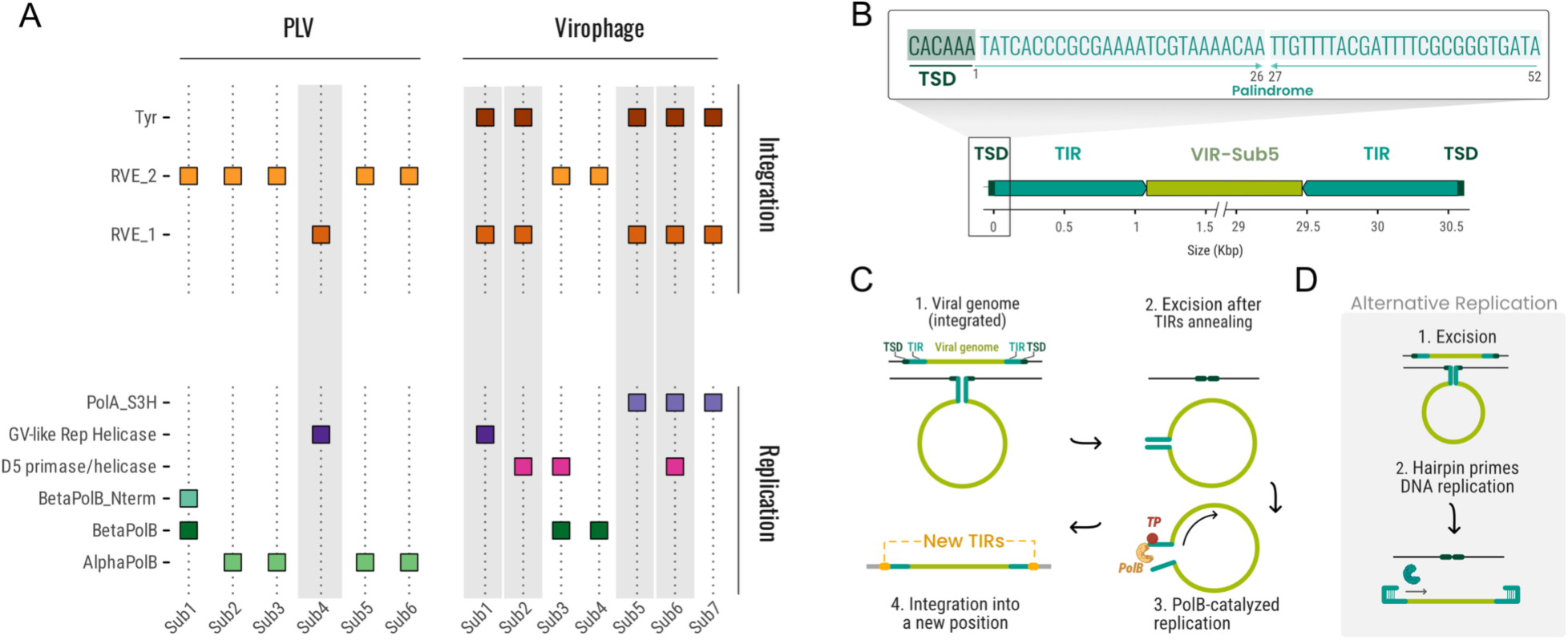
Integration and replication of *H. seosinensis* PLVs and virophages. **A.** Diverse assortment of genes associated with viral replication and integration in virophage and PLV subtypes. Subtypes containing terminal palindromic repeats are highlighted in light gray. **B**. Schematic representation of subtype 5 virophages, featuring 52-nucleotide palindromic sequences flanking the terminal inverted repeats (TIRs). **C**. Model of replication in which the viral genome is excised after annealing of opposing TIRs, replicated by pPolB, and integrated into a new genomic location marked by a new target site duplication (TSD) in orange. This model was proposed for polintons by Kapitonov et al. (3). **D**. Alternative model for viruses with flanking palindromic repeats. After excision, the palindromes at the TIR extremities form a hairpin which primes DNA replication. It may rely on a particular set of DNA replication enzymes.

Palindromic repeats at viral genome termini add another layer of potential mechanistic diversity. Systematic screening revealed palindromes within the TIRs and adjacent to TSDs (Fig. 4B) in all complete virophage subtypes 1, 2, 5, and 6, and in PLV subtype 4. These palindromes are preferentially associated with elements that use non–pPolB replication genes and particular RVE integrases (Fig. 4). Given that terminal hairpins prime replication in parvoviruses and poxviruses (39, 40), similar structures could contribute to replication of these endogenous elements in *H. seosinensis* (Fig. 4B). An alternative explanation is they may result from accidental DNA annealing caused by single-stranded DNA breaks (3). Experimental studies and a global characterization of these structures linked to replication/integration genes in *Polisviricotina* will be needed to confirm or refute this model.

### Dynamic evolution of virophages, PLVs and transposable elements

Endogenous virophages and PLVs in *H. seosinensis* frequently occur in nested arrangements, either within one another or within host transposable elements. We found multiple PLVs (n=5) and virophages (n=9) embedded inside other viral elements, including one striking case in which a virophage genome is both nested within a PLV and itself harbouring another virophage (Fig. 5A). In addition, PLVs and virophages are often inserted into regions annotated as repeats, including loci rich in Mutator-like DNA (MULE) transposons (Fig. 5B and Fig. S5A) and LINE retrotransposons (Fig. 5C and Fig. S5B). In some of these regions, transposase or reverse transcriptase genes were unidentifiable, suggesting they represent remnants of old transposons where viral sequences have integrated (Fig. S5C). These observations point to an active, multi-layered interplay between viral elements and host TEs.

**Fig. 5.**
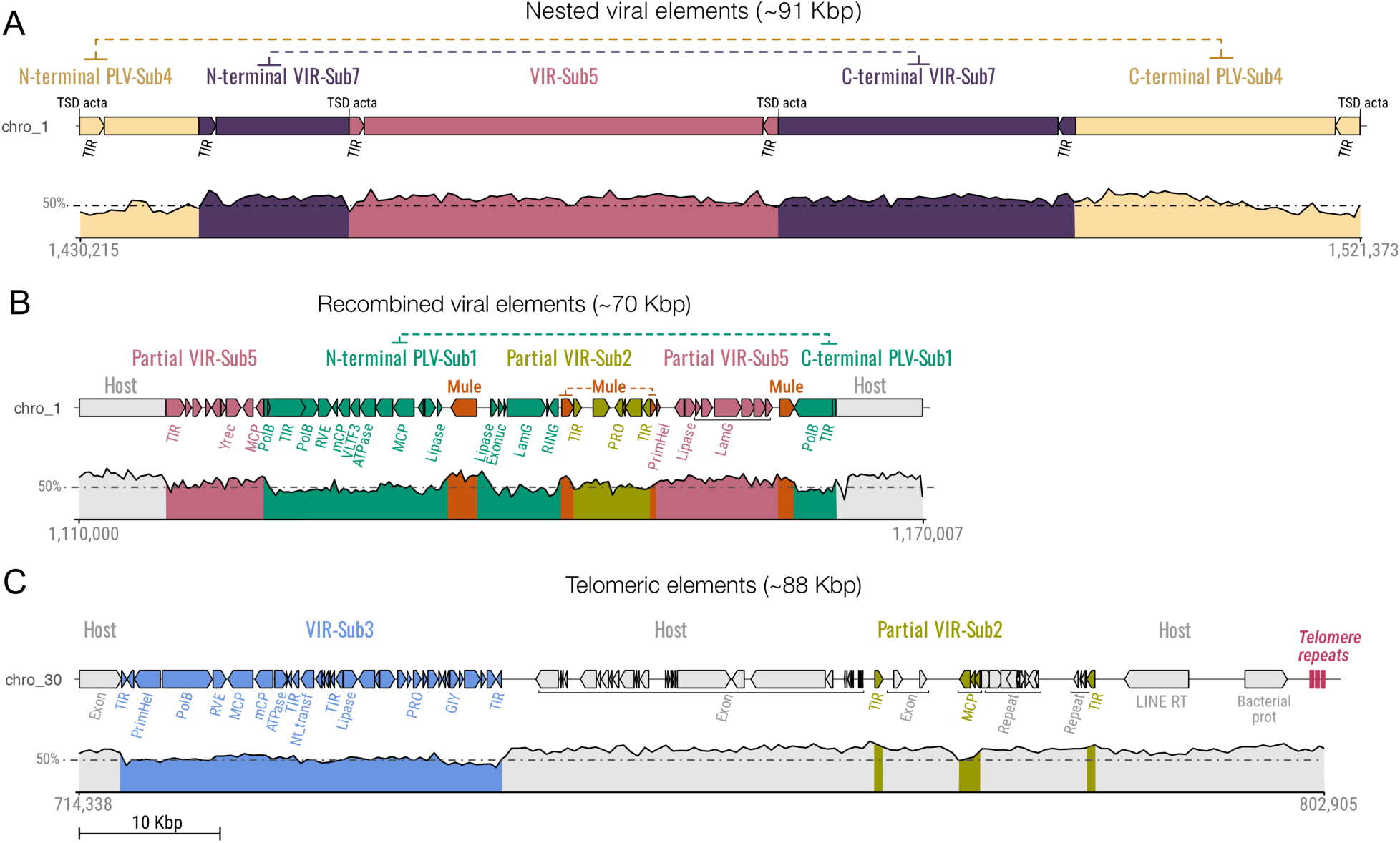
Genome dynamics of viral elements and host TEs in *H. seosinensis*. **A**. Supraparasitism of a host transposon (MULE) invading hypothetical recombined viral elements within a 60 Kbp genomic region on chromosome 1 (coordinates: 1,110,000-1,170,007). This region features partial virophage subtypes 5 and 2, a complete PLV subtype 1, and four copies of MULE transposon; GC content is colour-coded for each element and contrasted against the average host GC content of 47%. **B**. Nested arrangement of viral elements within a 91 Kbp region on chromosome 1 (coordinates: 1,430,215-1,521,373), displaying a complete virophage subtype 5 flanked by virophage subtype 7 and PLV subtype 4. Terminal inverted repeats (TIR) and target site duplications (TSD) are marked at element boundaries. **C**. Hotspot of viral and TE elements at the telomeric end of chromosome 30. A retrotransposon-derived (LINE) reverse transcriptase (RT) and a Deltaproteobacteria-like protein-coding gene were identified, along with virophage subtypes 2 and 3. Genomic representation and GC content estimation mirrors that shown in Fig. 1C.

TEs often integrate into PLVs or virophages genomes. The most common are MULEs, with 16 instances of insertion events detected in PLVs and virophages, identifiable through GC content variation (Fig. 5B, Fig. S6A-B). Both TIR-flanked and TIR-lacking forms were observed (Fig. S6A and Fig. S6B). Their distribution spans several viral subtypes and unrelated genomic regions, with outcomes ranging from apparent inactivation (e.g., disruption of a virophage replication gene, VIRO.tig00000004.961761.987735) to insertions that leave the core viral gene order intact (VIRO.tig00000065.137012.175777). In some genomic regions, full-length PLVs carry fragments of virophages from different subtypes together with multiple nearby MULE copies (Fig. 5B), consistent with TE-mediated recombination or gene shuffling among viral genomes. Other unclassified TEs, including elements with TIRs or reverse transcriptase genes adjacent to partial viral genomes lacking core genes, suggest that retrotransposition contributes to viral genome decay. Unlike *Cafeteria burkhardae* (21), *H. seosinensis* lacks Ngaro retrotransposons, indicating that similar supraparasitic dynamics can arise from different TE families in related hosts.

### Gene expression and methylation levels are low, giant-virus signatures are absent

Analysis of RNA-seq data revealed very low expression of genes in endogenous virophages and PLVs (Table S5), indicating that these elements were not actively replicating or transposing under the conditions sampled. In organisms where virophages and/or PLVs depend on co-infecting *Nucleocytoviricota* for replication (30), low expression of endogenous elements is interpreted as the absence of a giant virus partner. In searching the *H. seosinensis* genome, we did not find large blocks of viral genes as would be expected if there was a giant virus lurking in the system. We did, however, identify ∼250 predicted proteins with weak similarity to CroV paralogs of unknown function, CroV being the virus that infects the relative of *H. seosinensis*, *C. burkhardae*. A specific subclade of virophages in *H. seosinensis* also has promoters similar to a giant-viral promoter motif (Fig. S7A, Table S6). Such similarity seems to play a role in virophage reactivation upon giant virus infection, as shown in *C. burkardae* (14), and a similar mechanism may operate in *H.seosinensis*

To assess whether low expression might be linked to epigenetic silencing, we examined ONT sequence data for 5-methycytosine (5mC) methylation across viral regions within the *H. seosinensis* genome. We found no convincing evidence for localized spikes or hypermethylated motifs in the endogenized viral elements relative to the genomic background (Fig. S9), recognizing that small differences (Fig. S9B) can be artifacts of lower-quality base calls in low-coverage, compositionally biased, and homopolymer-rich regions (41, 42). We also found no genes encoding canonical eukaryotic methyltransferases (PF00145) in the *H. seosinensis* genome. There is thus no convincing evidence that 5mC methylation broadly suppresses PLVs or virophages in *H. seosinensis*, although other methylation modifications (e.g., 6mA and 4mC) could be involved. Low expression likely instead reflects the absence of appropriate activating conditions and/or the presence of decaying elements that are no longer capable of reactivation.

## DISCUSSION

By analyzing a long-read genome assembly of the hypersaline protist *H. seosinensis* (36), we have found dozens of PLVs and virophages integrated into the genome. In *C. burkardae*, the other bicosoecid with a sequenced genome, multiple endogenous virophages were previously described (21), but these insertions are not related to those of *H. seosinensis*. Some *C. burkardae* virophages are known to be associated with the *Nucleocytoviricota* CroV, the former providing immunity to the protist host against the latter (14, 43). To our knowledge, no member of the *Nucleocytoviricota* infecting *Halocafeteria* species has been identified. We were unable to find genomic signatures suggesting the existence of such a virus that could infect *H. seosinensis* and trigger the replication of some of its endogenous virophages or PLVs. Nonetheless, the presence of numerous CroV homologs in the *H. seosinensis* assembly, as well as virophage promoter motifs similar to the CroV ‘late’ motif suggests that a CroV-like giant virus could be, or could recently have been, associated with *H. seosinensis* and some of its endogenous viruses in nature. Note that a virophage unrelated to Mavirus was isolated together with a CroV-like *Nucleocytoviricota* in the green alga *Chlorella sp*. (44), showing that unrelated virophages can be associated with related *Nucleocytoviricota*.

Another difference between the *H. seosinensis* and *C. burkhardae* genomes is that while the former has both virophages and PLVs, the latter has only virophages. Our observation that in *H. seosinensis*, some homologs are found in subsets of PLVs and subsets of virophages — and that they are most closely related to each other in phylogenetic trees — is compatible with gene exchange between *H. seosinensis* endogenous virophages and PLVs. Other protists have both PLVs and virophages, e.g., *B. natans* and its distant cercozoan relative *Paulinella micropora* (18, 20). Their coexistence in *H. seosinensis* and other protists provides a common gene pool that supports the diversification of both viral-derived elements.

In *C. burkhardae*, cases of superparasitism in which endogenous virophages are themselves invaded by other mobile elements have been described, e.g., the tyrosine retrotransposon Ngaro landing in several integrated viral genomes (21). We have shown that *H. seosinensis* virophages and PLVs have also been invaded by transposable elements, in this case mainly by MULE DNA transposons and LINE retrotransposons (not Ngaro) — this represents another parallel yet distinct evolutionary path in the two bicosoecids. In some instances, the integrity of the viral genomes is apparently preserved despite MULE insertion. The transposon could thus conceivably be shuttled by the replicating viruses. In other cases, the TEs are in the vicinity of partial virus genomes, consistent with the hypothesis that TE integration triggers pseudogenization and turnover of viral genomes.

The presence of virophages, PLVs, transposable elements, and foreign DNA (such as a deltaproteobacterial gene (Fig. 5C)) in the subtelomeric regions of *H. seosinensis* pseudochromosomes suggests that these genomic regions could be a ‘safe harbour’ for viruses and TEs, as proposed in (45). Such regions typically lack essential host genes and thus could be more accessible for colonization. Viruses of other taxa, including herpesviruses (46), mirusviruses (47) and *Nucleocytoviricota* (45) have been found in the sub-telomeric regions of their eukaryotic host chromosomes (TEs are also overrepresented in these areas) (48–51). This appears to be the case for virophages and PLVs as well. Chase et al. (25) found that PLVs in the genomes of several green algae in the family *Chlorodendraceae* are not randomly distributed and may be localized to centromeres (25). Finally, we have shown that viral sequences are themselves hotspots for virophage and PLV insertions in the *H. seosinensis* genome, reminiscent of so-called twintrons (i.e., introns within introns (52)). The genomes of viruses and virus-like elements will land where they are tolerated; their persistence and spread have the potential to play an important role in shaping genome biology and evolution.

## MATERIAL AND METHODS

### Detection of polintons, polinton-like viruses, and virophages

We combined two approaches for the identification of viral regions in the *H. seosinensis* genome (36): (i) structural genes of known virophages and PLVs available in NCBI were used as probes in Blastp v.5 (53) and hmmsearch v.3.4 (54) similarity searches of the assembly; and (ii) regions exhibiting a reduced GC content were manually inspected using Geneious v.R10.0 (many endogenized viruses have GC contents that differ from their host genome (55)). Potential viral regions were further examined for the presence of TIRs using Blastn. Palindromic sequences within the viral genomes were detected using the EMBOSS palindrome tool (56) with the following parameters: -minpallen 20 - maxpallen 300 -nummismatches 4 -gaplimit 10 -overlap. ORFs were predicted using Prodigal v2.6.3 (57) with default parameters.

### Viral network reconstruction

Viral network analyses were conducted using several approaches. For the MCP network, PLV and virophage MCPs publicly available on NCBI (January 2024) were analyzed together with those identified in *H. seosinensis* using the Enzyme Similarity Tool (EFI-EST) (58) to create a sequence similarity network (SSN). A threshold score of 15 was chosen. The SSN was visualized (organic layout) and edited using Cytoscape v.3.10.3 (59). The viral proteome network was constructed based on the genomes and proteomes of virophages, polintons, and PLVs sourced from multiple studies (20, 22, 24, 25, 28, 37, 60, 61) using vConTACT2 v.0.11.3 (62) with the following parameters: -Diamond; - reported-alignments 20000; ClusterONE for VC, MCL for PC; -pc-inflation 1.8. Due to the high complexity of the resulting network, we extracted a subset of the data focused on *H. seosinensis* endogenous viruses and directly linked nodes for separate analysis. This subnetwork was visualized in Cytoscape v.3.10.3 using an edge-weighted spring-embedded model as described in (62).

### Protein cluster annotation

Protein clusters (PCs) were annotated by aligning each PC using MUSCLE. A Hidden Markov Model (HMM) profile was then built for each alignment using HMMER v.3.4 (54). These profiles were subsequently used to interrogate Swiss-Prot (https://www.uniprot.org/) databases (hmmsearch E-value cutoff 0.001). The top protein hit from each search was selected to create an annotation file. Manual annotation was performed using Blast and HHpred (63) to ensure accuracy for the most highly represented PCs. Genome visualization of representative viral elements and their predicted and annotated proteins was conducted using gggenomes v.0.9.5 (https://github.com/thackl/gggenomes) in R.

### Phylogenetic analysis

Amino acid sequence alignments were produced using MAFFT v7.490 (64) with dataset-specific parameters. Alignments were typically refined using TrimAl v.1.4 (65) for site selection. Phylogenetic trees were constructed using IQ-TREE v2.0.7 (66), employing an automated model selection and 1,000 ultrafast bootstrap replicates.

### Promoter motif analysis

For each viral genome, putative promoter motifs were identified by analyzing the first 100 nucleotides upstream of each ORF predicted by prodigal. These motifs were then analyzed using MEME v.5.5.5 (67) to detect the best-scoring motif per element. Motif search parameters were set to identify sequences 4 to 10 nucleotides in length, with an E-value threshold of less than 0.1, and zero to one motif per sequence. The inferred promoter motifs were used as queries to search for similar motifs in other elements.

### Genome synteny analysis

To visualize the nucleotide percent identity among virophages and PLVs within each subtype, the online tool DiGAlign (68) was employed. The parameters ‘BLASTn’ and ‘Without gene information’ were selected.

### Gene expression and methylation

Six previously generated RNA-seq datasets (35) were utilized to estimate the expression of viral regions and host genes in *H. seosinensis* (accession numbers SRX1423011, SRX1423005, SRX1423004, SRX1421934, SRX1423015, and SRX1423016). RNA-seq reads were mapped to the new genome assembly using HISAT2 v.2.2.1 (69). The software featureCounts v.1.5.2 (70) was used to quantify expression. Raw read counts for each feature (gene) were divided by gene length for normalization. Guppy (v6.5.7) was used to detect 5mC bases across the genome, using the Rerio all-contexts modified base-calling model res_dna_r941_min_modbases-all-context_v001. Methylation-called reads were then mapped onto the assembly using Minimap2, and the methylation levels at each site were calculated using modbam2bed.

## ACKNOWLEDGEMENTS

Yana Eglit is thanked for contributing *Halocafeteria seosinensis* cultures, and Guillaume Blanc, Tommy Harding and Sebastian Hess are thanked for early insights on the analyses carried out herein.

## REFERENCES

1. J. N. Wells, C. Feschotte, A Field Guide to Eukaryotic Transposable Elements. Annu. Rev. Genet. 54, 539–561 (2020).

2. C. Feschotte, E. J. Pritham, DNA Transposons and the Evolution of Eukaryotic Genomes. Annu. Rev. Genet. 41, 331–368 (2007).

3. V. V. Kapitonov, J. Jurka, Self-synthesizing DNA transposons in eukaryotes. Proc. Natl. Acad. Sci. 103, 4540–4545 (2006).

4. M. Krupovic, D. H. Bamford, E. V. Koonin, Conservation of major and minor jelly-roll capsid proteins in Polinton (Maverick) transposons suggests that they are bona fide viruses. Biol. Direct 9, 6 (2014).

5. M. G. Fischer, The Virophage Family *Lavidaviridae*. Curr. Issues Mol. Biol. 40, 1–24 (2021).

6. S. Mougari, D. Sahmi-Bounsiar, A. Levasseur, P. Colson, B. La Scola, Virophages of Giant Viruses: An Update at Eleven. Viruses 11, 733 (2019).

7. S. Duponchel, M. G. Fischer, Viva lavidaviruses! Five features of virophages that parasitize giant DNA viruses. PLoS Pathog. 15, e1007592 (2019).

8. E. V. Koonin, M. Krupovic, N. Yutin, Evolution of double-stranded DNA viruses of eukaryotes: from bacteriophages to transposons to giant viruses. Ann. N. Y. Acad. Sci. 1341, 10–24 (2015).

9. M. G. Fischer, C. A. Suttle, A virophage at the origin of large DNA transposons. Science 332, 231–234 (2011).

10. B. La Scola, et al., The virophage as a unique parasite of the giant mimivirus. Nature 455, 100–104 (2008).

11. M. Boughalmi, et al., High-throughput isolation of giant viruses of the *Mimiviridae* and *Marseilleviridae* families in the Tunisian environment. Environ. Microbiol. 15, 2000–2007 (2013).

12. R. K. Campos, et al., Samba virus: a novel mimivirus from a giant rain forest, the Brazilian Amazon. Virol. J. 11, 95 (2014).

13. M. Gaia, et al., Zamilon, a novel virophage with *Mimiviridae* host specificity. PloS One 9, e94923 (2014).

14. A. Koslová, et al., Endogenous virophages are active and mitigate giant virus infection in the marine protist *Cafeteria burkhardae*. Proc. Natl. Acad. Sci. U. S. A. 121, e2314606121 (2024).

15. B. P. Taylor, M. H. Cortez, J. S. Weitz, The virus of my virus is my friend: ecological effects of virophage with alternative modes of coinfection. J. Theor. Biol. 354, 124– 136 (2014).

16. S. Yau, et al., Virophage control of antarctic algal host–virus dynamics. Proc. Natl. Acad. Sci. 108, 6163–6168 (2011).

17. J. G. Nino Barreat, A. Katzourakis, Ecological and evolutionary dynamics of cell-virus-virophage systems. PLoS Comput. Biol. 20, e1010925 (2024).

18. G. Blanc, L. Gallot-Lavallée, F. Maumus, Provirophages in the *Bigelowiella* genome bear testimony to past encounters with giant viruses. Proc. Natl. Acad. Sci. 112, E5318–E5326 (2015).

19. Y. Gyaltshen, et al., Long-Read-Based Genome Assembly Reveals Numerous Endogenous Viral Elements in the Green Algal Bacterivore *Cymbomonas tetramitiformis*. Genome Biol. Evol. 15, evad194 (2023).

20. C. Bellas, et al., Large-scale invasion of unicellular eukaryotic genomes by integrating DNA viruses. Proc. Natl. Acad. Sci. 120, e2300465120 (2023).

21. T. Hackl, S. Duponchel, K. Barenhoff, A. Weinmann, M. G. Fischer, Virophages and retrotransposons colonize the genomes of a heterotrophic flagellate. eLife 10, e72674 (2021).

22. N. Yutin, S. Shevchenko, V. Kapitonov, M. Krupovic, E. V. Koonin, A novel group of diverse Polinton-like viruses discovered by metagenome analysis. BMC Biol. 13, 95 (2015).

23. N. Yutin, D. Raoult, E. V. Koonin, Virophages, polintons, and transpovirons: a complex evolutionary network of diverse selfish genetic elements with different reproduction strategies. Virol. J. 10, 158 (2013).

24. C. M. Bellas, R. Sommaruga, Polinton-like viruses are abundant in aquatic ecosystems. Microbiome 9, 13 (2021).

25. E. E. Chase, C. Desnues, G. Blanc, Integrated viral elements suggest the dual lifestyle of *Tetraselmis spp*. Polinton-like viruses. Virus Evol. 8, veac068 (2022).

26. Z. K. Barth, et al., Genomic analysis of hyperparasitic viruses associated with entomopoxviruses. Virus Evol. 10, veae051 (2024).

27. M. Krupovic, J. H. Kuhn, M. G. Fischer, E. V. Koonin, Natural history of eukaryotic DNA viruses with double jelly-roll major capsid proteins. Proc. Natl. Acad. Sci. 121, e2405771121 (2024).

28. N. Yutin, V. V. Kapitonov, E. V. Koonin, A new family of hybrid virophages from an animal gut metagenome. Biol. Direct 10, 19 (2015).

29. S. Santini, et al., Genome of *Phaeocystis globosa* virus PgV-16T highlights the common ancestry of the largest known DNA viruses infecting eukaryotes. Proc. Natl. Acad. Sci. 110, 10800–10805 (2013).

30. S. Roitman, et al., Isolation and infection cycle of a polinton-like virus virophage in an abundant marine alga. Nat. Microbiol. 8, 332–346 (2023).

31. A. Pagarete, T. Grébert, O. Stepanova, R.-A. Sandaa, G. Bratbak, Tsv-N1: A Novel DNA Algal Virus that Infects *Tetraselmis striata*. Viruses 7, 3937–3953 (2015).

32. E. V. Koonin, M. G. Fischer, J. H. Kuhn, M. Krupovic, The polinton-like supergroup of viruses: evolution, molecular biology, and taxonomy. Microbiol. Mol. Biol. Rev. 0, e00086–23 (2024).

33. M. D. Johnson, G. M. Housey, P. T. Kirschmeier, I. B. Weinstein, Molecular cloning of gene sequences regulated by tumor promoters and mitogens through protein kinase C. Mol. Cell. Biol. 7, 2821–2829 (1987).

34. L. A. Sarre, et al., DNA methylation enables recurrent endogenization of giant viruses in an animal relative. Sci. Adv. 10, eado6406 (2024).

35. T. Harding, M. W. Brown, A. G. B. Simpson, A. J. Roger, Osmoadaptative Strategy and Its Molecular Signature in Obligately Halophilic Heterotrophic Protists. Genome Biol. Evol. 8, 2241–2258 (2016).

36. L. Gallot-Lavallée, et al., Sequencing, Chromosome-scale Assembly, and Annotation of the Genome of the Halophilic Nanoflagellate *Halocafeteria seosinensis*. [Preprint] (2026). Available at: https://www.biorxiv.org/content/10.64898/2026.06.25.734631v1 [Accessed 22 July 2026].

37. S. Roux, et al., Updated Virophage Taxonomy and Distinction from Polinton-like Viruses. Biomolecules 13, 204 (2023).

38. C. Desnues, et al., Provirophages and transpovirons as the diverse mobilome of giant viruses. Proc. Natl. Acad. Sci. U. S. A. 109, 18078–18083 (2012).

39. B. Moss, Poxvirus DNA replication. Cold Spring Harb. Perspect. Biol. 5, a010199 (2013).

40. F. S. De Silva, W. Lewis, P. Berglund, E. V. Koonin, B. Moss, Poxvirus DNA primase. Proc. Natl. Acad. Sci. U. S. A. 104, 18724–18729 (2007).

41. C. Delahaye, J. Nicolas, Sequencing DNA with nanopores: Troubles and biases. PLOS ONE 16, e0257521 (2021).

42. B. A. Curtis, et al., Algal genomes reveal evolutionary mosaicism and the fate of nucleomorphs. Nature 492, 59–65 (2012).

43. M. G. Fischer, T. Hackl, Host genome integration and giant virus-induced reactivation of the virophage mavirus. Nature 540, 288–291 (2016).

44. Y. Sheng, Z. Wu, S. Xu, Y. Wang, Isolation and Identification of a Large Green Alga Virus (Chlorella Virus XW01) of *Mimiviridae* and Its Virophage (Chlorella Virus Virophage SW01) by Using Unicellular Green Algal Cultures. J. Virol. 96, e0211421 (2022).

45. C. Blais, M. J. Colp, L. A. Sarre, A. de Mendoza, J. M. Archibald, Epigenetic silencing and genome dynamics determine the fate of giant virus endogenizations in *Acanthamoeba*. BMC Biol. 23, 171 (2025).

46. N. Osterrieder, N. Wallaschek, B. B. Kaufer, Herpesvirus Genome Integration into Telomeric Repeats of Host Cell Chromosomes. Annu. Rev. Virol. 1, 215–235 (2014).

47. J. L. Collier, et al., The protist *Aurantiochytrium* has universal subtelomeric rDNAs and is a host for mirusviruses. Curr. Biol. CB 33, 5199–5207.e4 (2023).

48. G. Servant, P. L. Deininger, Insertion of Retrotransposons at Chromosome Ends: Adaptive Response to Chromosome Maintenance. Front. Genet. 6, 358 (2015).

49. F. Chaux-Jukic, et al., Architecture and evolution of subtelomeres in the unicellular green alga *Chlamydomonas reinhardtii*. Nucleic Acids Res. 49, 7571–7587 (2021).

50. N. W. G. Chen, et al., Common Bean Subtelomeres Are Hot Spots of Recombination and Favor Resistance Gene Evolution. Front. Plant Sci. 9, 1185 (2018).

51. E. Fabre, et al., Comparative genomics in hemiascomycete yeasts: evolution of sex, silencing, and subtelomeres. Mol. Biol. Evol. 22, 856–873 (2005).

52. R. G. Drager, R. B. Hallick, A complex twintron is excised as four individual introns. Nucleic Acids Res. 21, 2389–2394 (1993).

53. C. Camacho, et al., BLAST+: architecture and applications. BMC Bioinformatics 10, 421 (2009).

54. S. R. Eddy, Accelerated Profile HMM Searches. PLoS Comput. Biol. 7, e1002195 (2011).

55. T. Hackl, et al., Four high-quality draft genome assemblies of the marine heterotrophic nanoflagellate *Cafeteria roenbergensis*. Sci. Data 7, 29 (2020).

56. P. Rice, I. Longden, A. Bleasby, EMBOSS: The European Molecular Biology Open Software Suite. Trends Genet. 16, 276–277 (2000).

57. D. Hyatt, et al., Prodigal: prokaryotic gene recognition and translation initiation site identification. BMC Bioinformatics 11, 119 (2010).

58. R. Zallot, N. Oberg, J. A. Gerlt, The EFI Web Resource for Genomic Enzymology Tools: Leveraging Protein, Genome, and Metagenome Databases to Discover Novel Enzymes and Metabolic Pathways. Biochemistry 58, 4169–4182 (2019).

59. P. Shannon, et al., Cytoscape: a software environment for integrated models of biomolecular interaction networks. Genome Res. 13, 2498–2504 (2003).

60. S. Roux, et al., Ecogenomics of virophages and their giant virus hosts assessed through time series metagenomics. Nat. Commun. 8, 858 (2017).

61. G. J. Starrett, et al., Adintoviruses: a proposed animal-tropic family of midsize eukaryotic linear dsDNA (MELD) viruses. Virus Evol. 7, veaa055 (2021).

62. H. Bin Jang, et al., Taxonomic assignment of uncultivated prokaryotic virus genomes is enabled by gene-sharing networks. Nat. Biotechnol. 37, 632–639 (2019).

63. J. Söding, A. Biegert, A. N. Lupas, The HHpred interactive server for protein homology detection and structure prediction. Nucleic Acids Res. 33, W244–W248 (2005).

64. K. Katoh, D. M. Standley, MAFFT Multiple Sequence Alignment Software Version 7: Improvements in Performance and Usability. Mol. Biol. Evol. 30, 772–780 (2013).

65. S. Capella-Gutiérrez, J. M. Silla-Martínez, T. Gabaldón, trimAl: a tool for automated alignment trimming in large-scale phylogenetic analyses. Bioinformatics 25, 1972– 1973 (2009).

66. L.-T. Nguyen, H. A. Schmidt, A. von Haeseler, B. Q. Minh, IQ-TREE: A Fast and Effective Stochastic Algorithm for Estimating Maximum-Likelihood Phylogenies. Mol. Biol. Evol. 32, 268–274 (2015).

67. T. L. Bailey, C. Elkan, Fitting a mixture model by expectation maximization to discover motifs in biopolymers. Proc. Int. Conf. Intell. Syst. Mol. Biol. 2, 28–36 (1994).

68. Y. Nishimura, K. Yamada, Y. Okazaki, H. Ogata, DiGAlign: Versatile and Interactive Visualization of Sequence Alignment for Comparative Genomics. Microbes Environ. 39, ME23061 (2024).

69. D. Kim, B. Langmead, S. L. Salzberg, HISAT: a fast spliced aligner with low memory requirements. Nat. Methods 12, 357–360 (2015).

70. Y. Liao, G. K. Smyth, W. Shi, featureCounts: an efficient general purpose program for assigning sequence reads to genomic features. Bioinformatics 30, 923–930 (2014).

71. R. C. Edgar, Search and clustering orders of magnitude faster than BLAST. Bioinforma. Oxf. Engl. 26, 2460–2461 (2010).

