## Supplementary data for "Multiple origins of endogenous virophage and polinton-like virus in the halophilic protist *Halocafeteria seosinensis*"

### SUPPLEMENTARY FIGURES

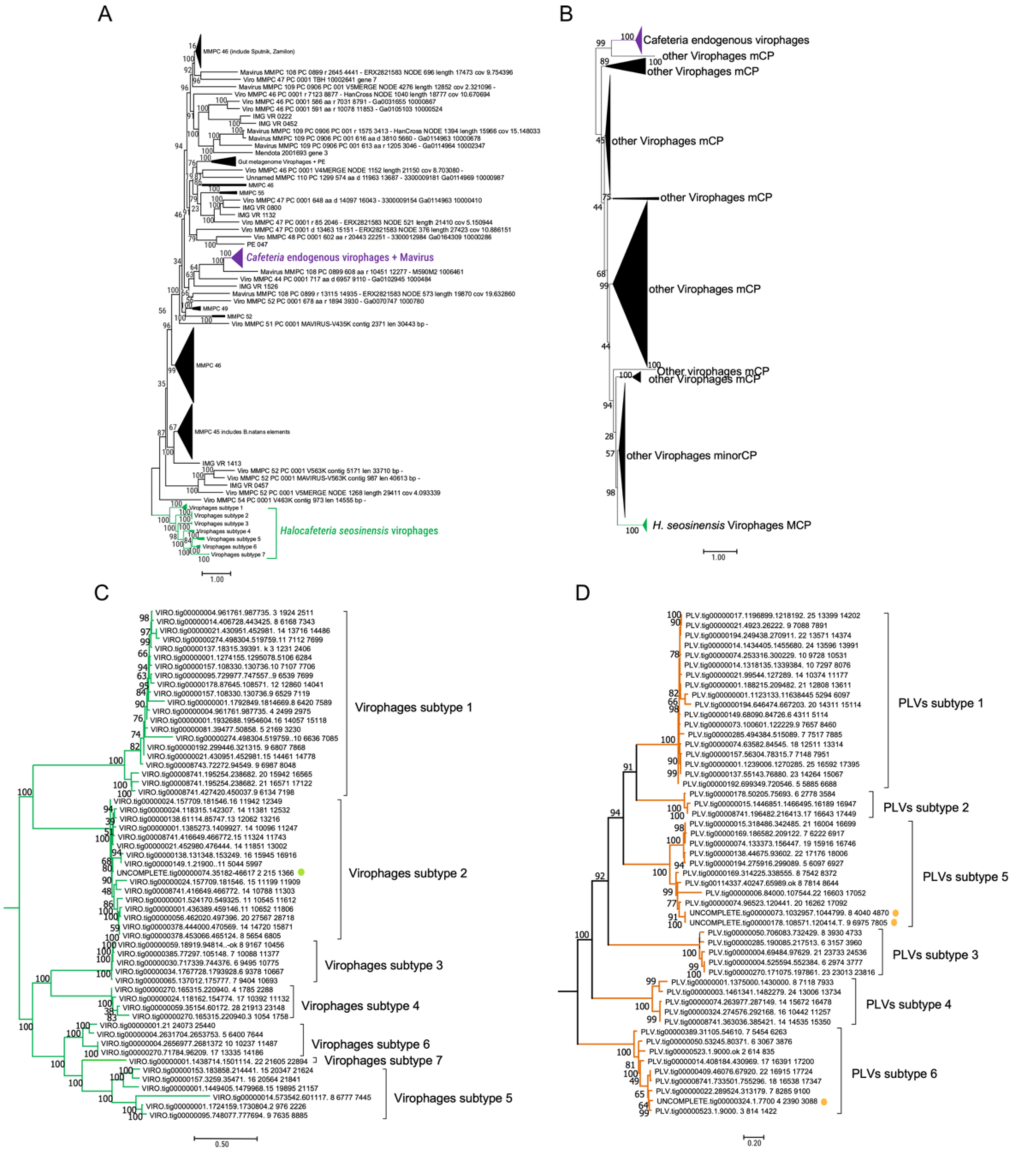


**Fig. S1. Phylogenetic reconstruction of major and minor capsid proteins in virophages from *H. seosinensis*. A**. Maximum likelihood (ML) phylogeny of virophage MCP (LG+F+R10 model and 1000 ufr). 629 sequences were aligned with mafft-linsi, and 510 sites with <30% gaps were retained. **B**. Virophage ML mCP phylogeny (LG+F+R10 model and 1000 ufb). 2,060 sequences were aligned with mafft-linsi, and 350 sites with <30% gaps were retained. **C**. Subtree containing *H. seosinensis* endogenous virophage mCPs of section B. **D**. ML phylogeny of mCP from PLVs (VT+F+R6 model and 1000 ufb). 421 sequences were aligned with mafft-linsi, and 280 sites with <30% gaps were retained. The subtree containing *H. seosinensis* endogenous PLV mCPs was extracted.

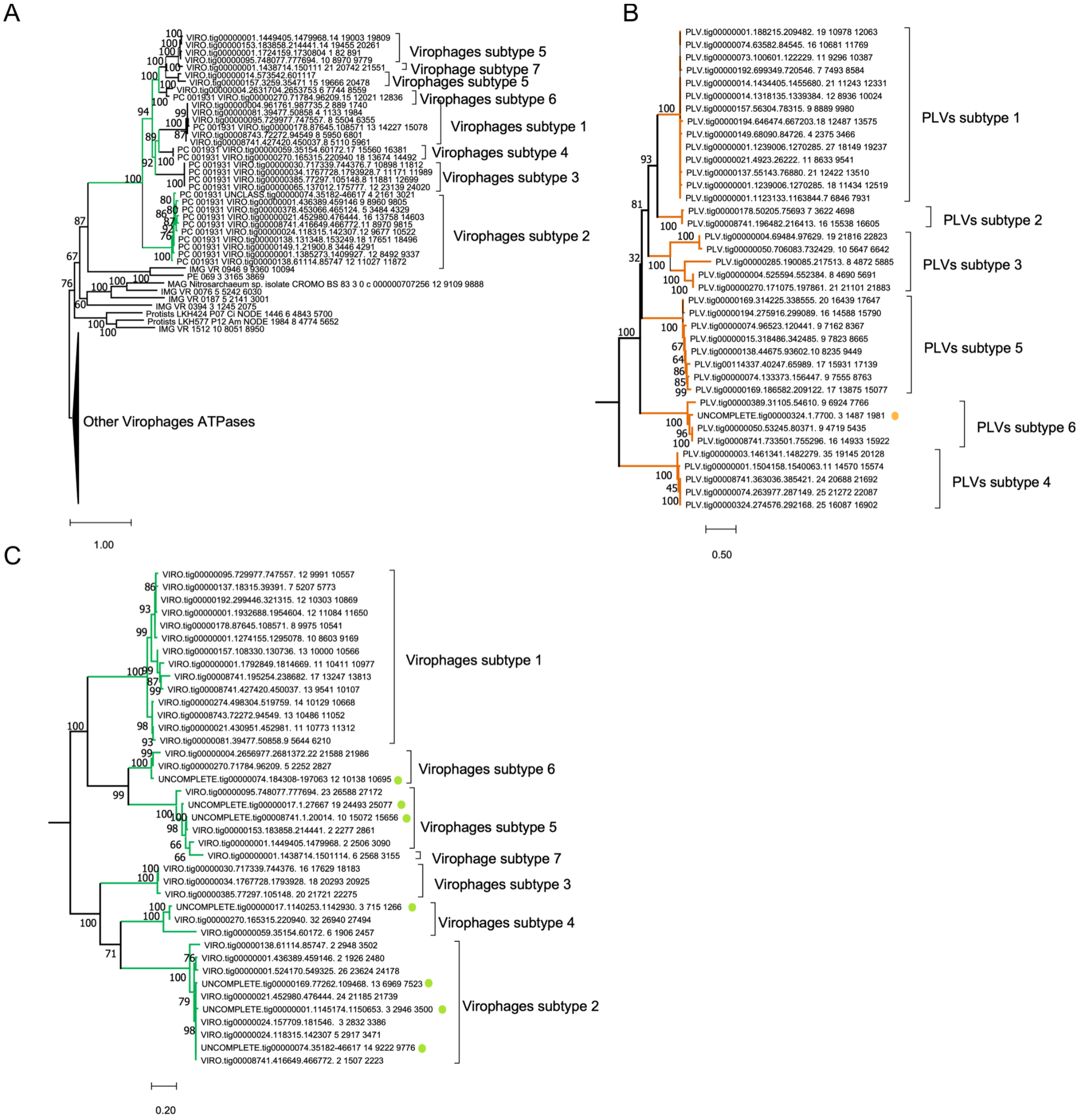


**Fig. S2. ML phylogenetic reconstruction of ATPase and PRO**. **A**. Virophage ATPase tree inferred under the LG+F+R8 model and 1000 ultrafast bootstrap. 271 sequences were aligned with mafft-linsi; 242 sites with <30% gaps were retained. **B**. PLV ATPase tree inferred under the LG+F+R5 model and 1000 ultrafast bootstrap. 129 sequences were aligned with mafft-linsi, with 226 sites having <30% gaps being retained. The subtree containing *H. seosinensis* PLV ATPases was extracted. **C**. Virophage PRO tree inferred under the LG+F+R5 model and 1000 ultrafast bootstrap. 147 sequences were aligned with mafft-linsi, and 168 sites with <30% gaps were retained. The subtree containing *H. seosinensis* virophages PRO was extracted.

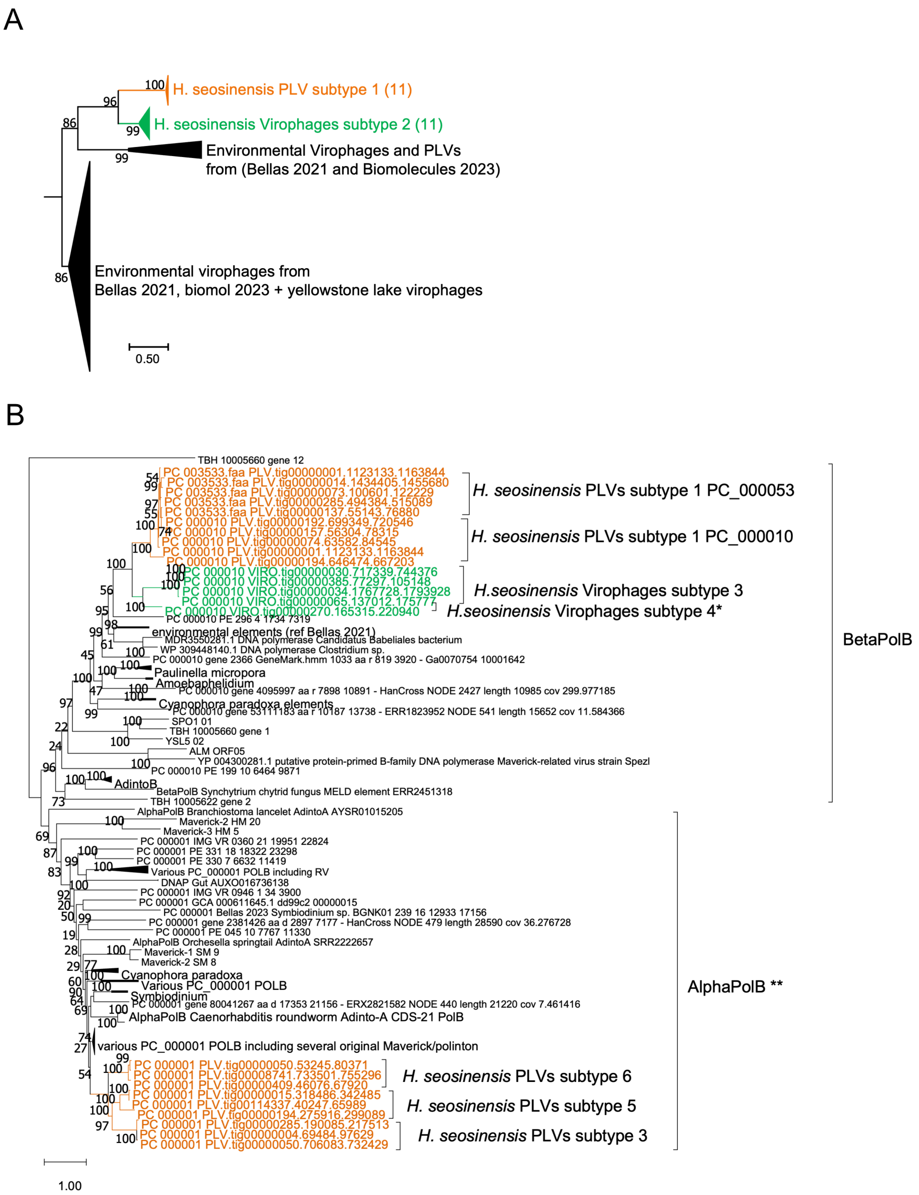


**Fig. S3. Gene exchange between *H. seosinensis* PLVs and virophages**. **A**. RING domain ML phylogeny (LG+I+G4 1000 ufb). 152 sequences were aligned with mafft-linsi, and 121 sites with <30% gaps were retained. **B**. pPolB ML phylogeny (LG+F+R6 model and 1000 ufb). A subset of sequences from PC_000001, PC_000010 and PC_000053 were chosen using uclust. 141 final sequences were aligned with mafft-linsi, with 836 sites containing <30% gaps being retained. The N-terminus of pPolB was duplicated in PLVs subtype 1 (PC_000053). **Sequences from the subtype 2 PLVs and subtype 4 were removed due to their divergent nature and poor alignment.

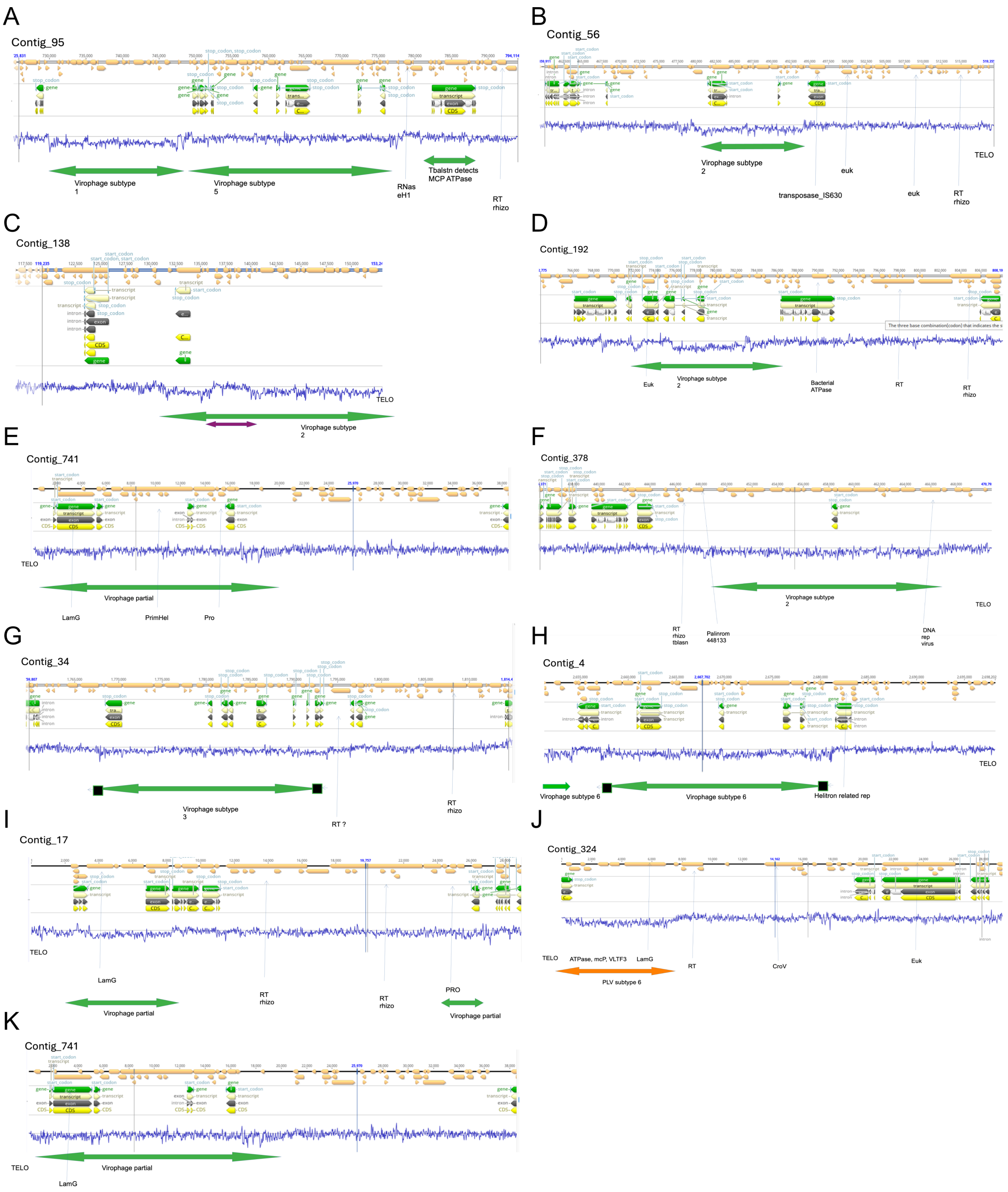


**Fig. S4. Endogenous virophages and PLVs at telomeres of *H. seosinensis* pseudochromosomes (A-K).** Screenshots of *H. seosinensis* telomeres displaying enrichment in these elements. These genomic regions were roughly annotated using Geneious v.R10.

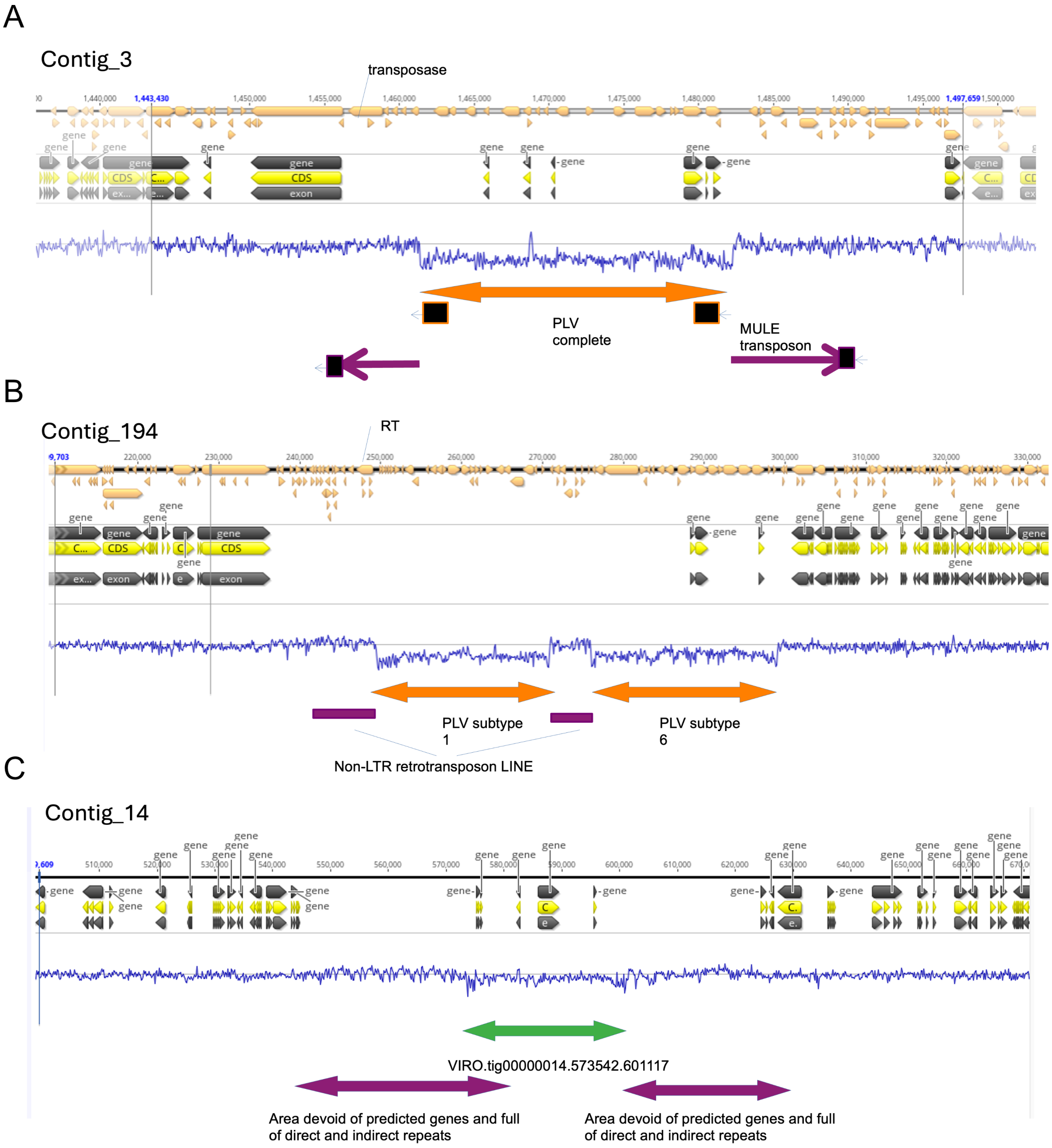


**Fig. S5. Endogenous virophages and PLVs were found inserted in host mobile elements and repeats**. **A**. PLV genome located within a Mule transposon. **B**. A PLV genome residing within a LINE retrotransposon. **C**. A virophage genome full of repeats on *H. seosinensis* contig 14.

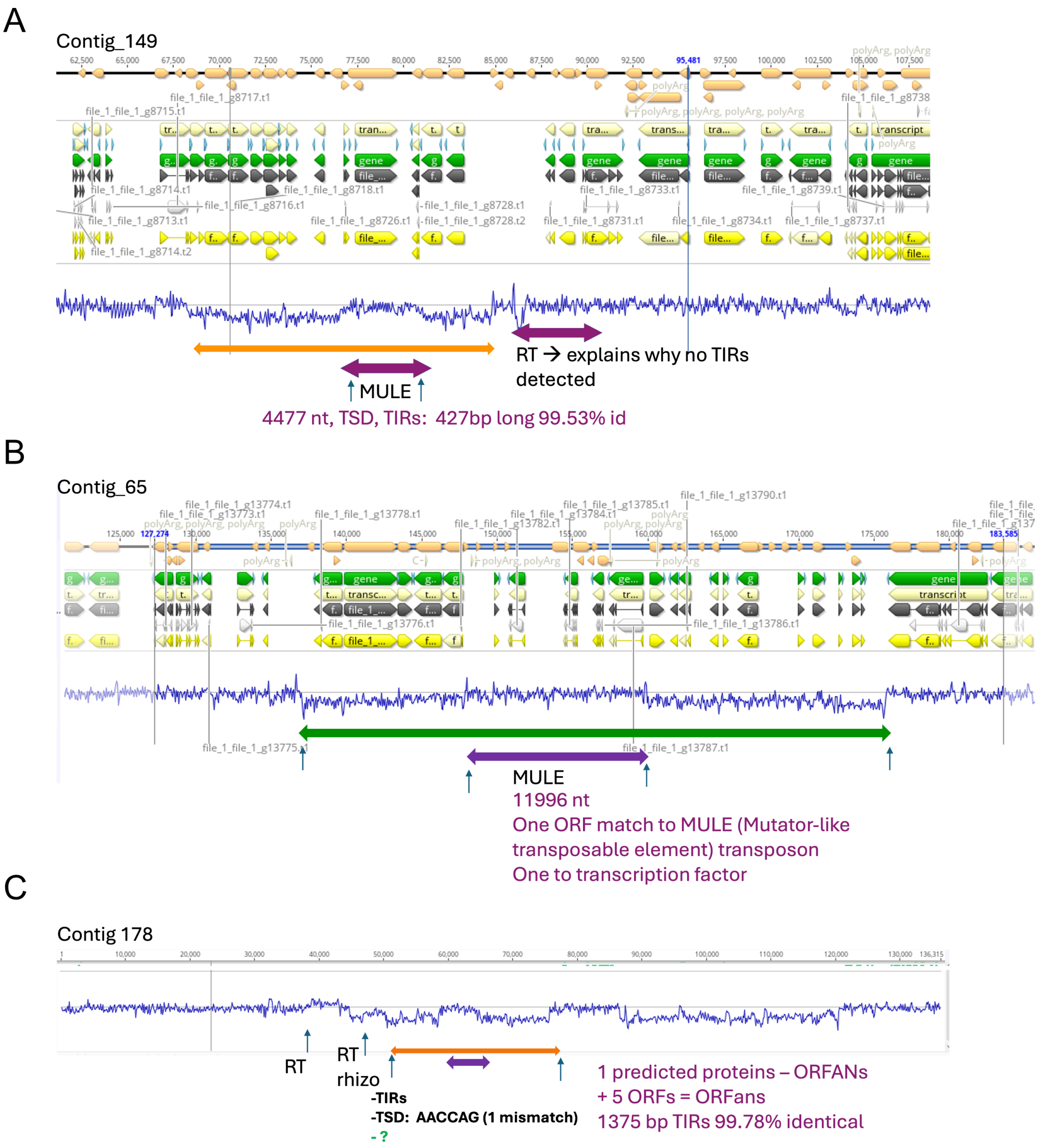


**Fig. S6. Supraparasitism of *H. seosinensis* endogenous viruses**. **A**. The PLV genome in contig 149 harbours a MULE transposon and a retrotransposon. No TIRs were detected. **B**. A transposon containing a MULE transposase is integrated within a virophage genome. No TIRs were detected around the transposase. The transposon does not appear to have interrupted any virophage genes. The viral genome contains all core genes, TIRs, and TSDs. **C**. An unclassified TE within a PLV genome; collinearity of the PLV with a related transposon-lacking PLV is apparent.

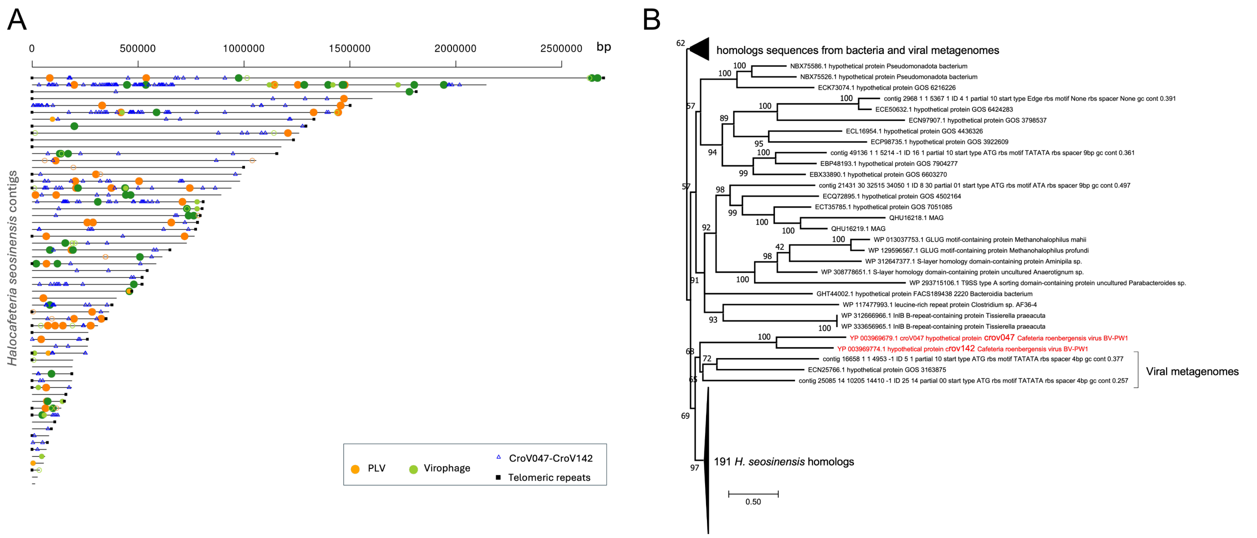


**Fig. S7. Traces of *Nucleocytoviricota* in the nuclear genome of *H. seosinensis*. A.** Location of the endogenous viruses and predicted proteins homologous to CroV047 and CroV142 in the *H. seosinensis* assembly. **B**. Phylogenetic tree of CroV047 and CroV142 homologs inferred under the LG+R8 model and 1000 ultrafast bootstraps. Sequences longer than 250 amino acids were aligned with MAFFT. 379 sites were manually selected. *Cafeteria roenbergensis*-associated *Nucleocytoviricota* CroV are in red.

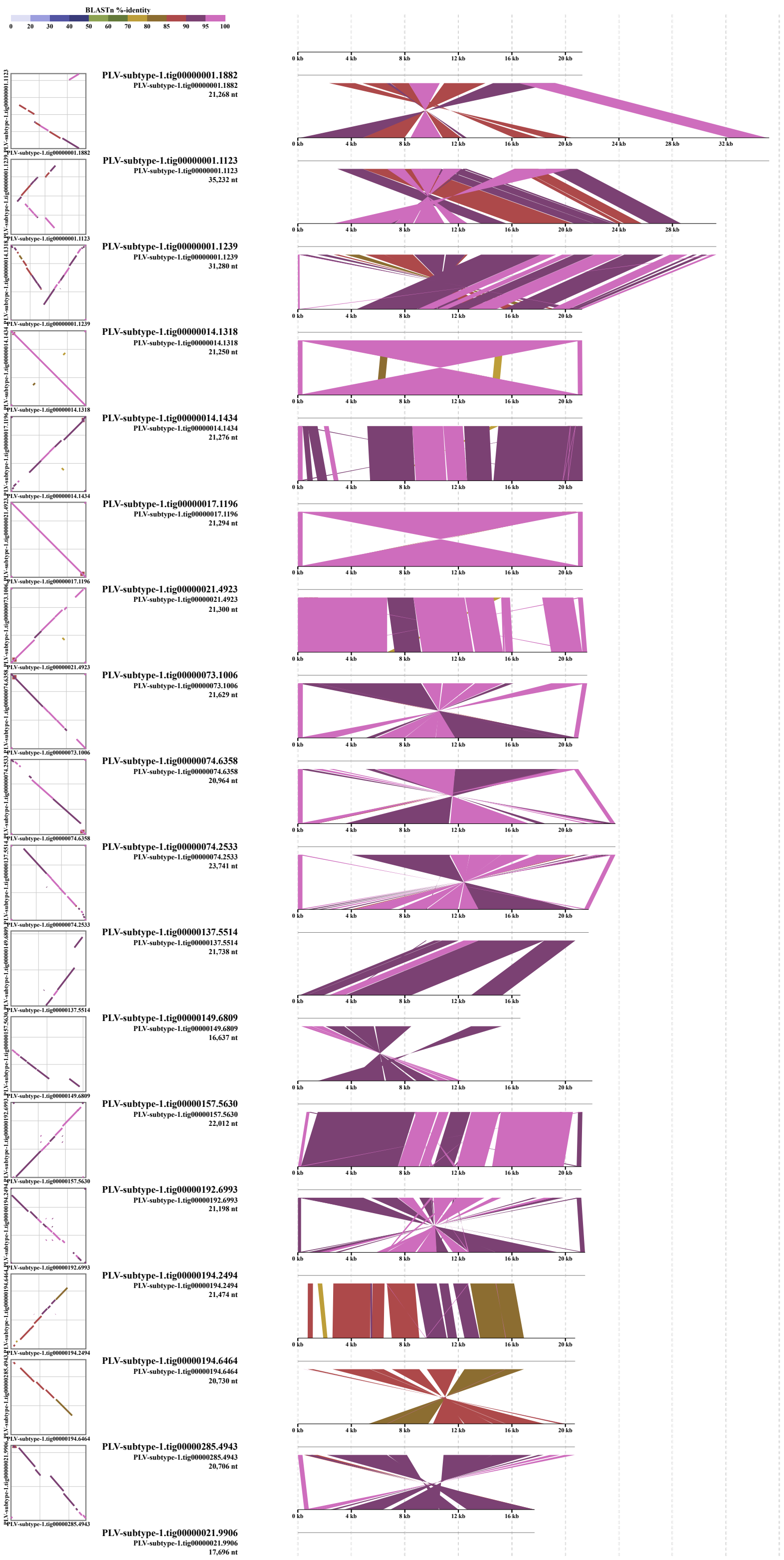


**Fig. S8. Genome alignment of subtype 1 PLVs from *H. seosinensis***. Additional PLVs and alignments are provided in supplementary files.

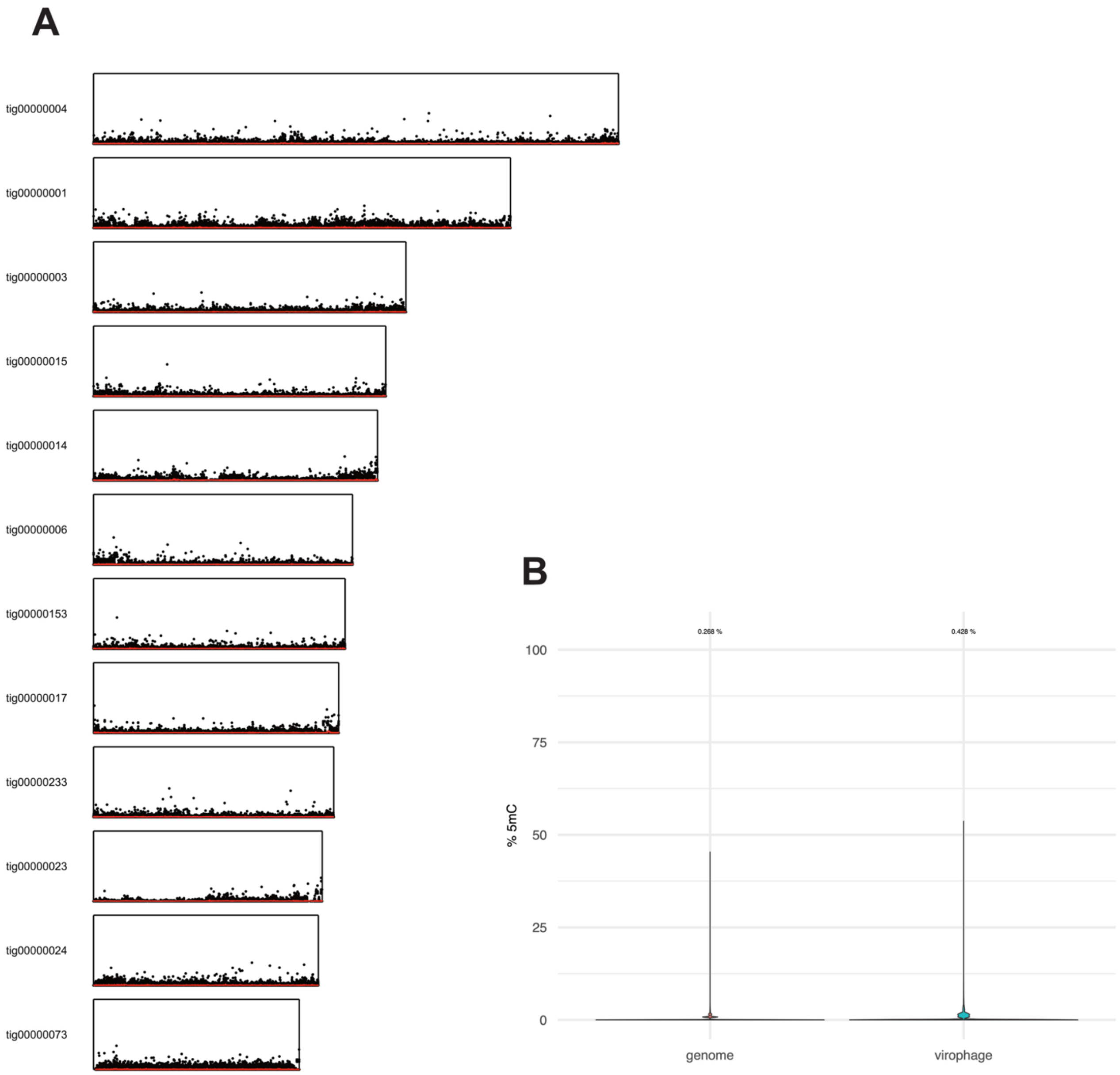


**Fig. S9. Lack of evidence for methylation in the *H. seosinensis* nuclear genome. A**. Global map of 5mC methylation levels across all contigs larger than 1 Mbp. Each methylation-called cytosine is shown as a single point along a contig. The position of the point on the y-axis corresponds to the percentage of reads methylated at that site (min 0%, max 100%). The rolling mean of methylation is shown in red. Only sites with a read coverage greater than 50 were included. **B**. Violin plot showing the distribution of methylation levels across all cytosines in virophages compared to the rest of the genome.

Only sites with a read coverage greater than 50 were included in this analysis. Most values in both violin plots are zero.
